# A Subtle Balance between Host Selection and Microbial Size Effect Mediates Plant Microbiome Assembly in Mulberry

**DOI:** 10.1101/2022.09.14.507911

**Authors:** Jintao He, Xiaoqiang Shen, Nan Zhang, Abrar Muhammad, Yongqi Shao

## Abstract

Breeding toward improved ecological plant–microbiome interactions requires improved knowledge of ecological processes/principles underlying microecological patterns, but these remain elusive. Here, we systematically investigated *in-planta* biogeographic patterns of plant-associated bacteriome and mycobiome along soil– plant and episphere–endosphere continuum in *Morus alba*. Microecological patterns in diversity, structure, co-occurrence network, species turnover, and assembly process were revealed and varying between bacteriome and mycobiome, possibly driven by multiple factors including host selection, community cohesion, and particularly size-dependent dispersal limitation. Based on these findings, we hypothesized that host selection historically recruits microbiotas, whereas microbial size affects the subsequent turnovers due to the limited dispersal of larger-size microbes. This hypothesis was supported by datasets from other plant species and confirmed by stochastic dispersal experiments showing that smaller-size microbes are more likely to escape/disperse from endosphere niches, contributing to fleeting niches occupied by larger-size microbes. These findings may open new avenues toward an improved understanding of the dynamics of plant microbiome assembly.

## 1 Introduction

Plants harbour numerous and diverse prokaryotes and eukaryotes that are well adapted to life on and inside plant tissues [1]. Although plants have evolved their adaptations to cope with various biotic and abiotic stresses, they also rely on their microbial partners [2-4]. Plant microbiota acts as a highly diversified, external secondary genome for the host and provides several essential functionalities [5], including disease suppression [6], nutrient acquisition [7], and adaptation to environmental variations [8]. In return, the host plant provides habitats and constant supplies of energy and carbon sources to the microbiota [6]. A current concept considers the plant and its associated microbiota as a functional entity called “holobiont” [9]. Thus, targeted manipulation of plant microbiomes offers a promising and sustainable approach for maximizing plant production to meet the challenges of global change. Of course, it requires that we understand the principle governing the complex assemblage of plant microbiomes.

Plant-associated microbiotas inhabit multiple microhabitats (i.e., niches) along the soil–plant continuum (SPC) and are compartmentalized by both external (episphere) and internal tissues (endosphere). Several studies have highlighted that plant microbiotas are tightly controlled by an intensive process [10, 11]. Significant and major host effects have been reported in crop microbiomes and the selection pressure on the bacterial community increases along the SPC [12]. Apart from host effects, growing evidence suggest that microorganisms themselves do follow some classic ecological patterns and mechanisms (e.g., species turnover, taxon co-occurrence, community cohesion, and prior effect) [13]. For instance, co-occurrence analysis of the oral microbiota showed significant segregation amongst bacterial taxa when community membership was examined at the level of genus but not at the level of species, suggesting that ecologically significant, competitive interactions are more apparent at a broader taxonomic level than species [14]. It has been reported in the aquatic ecosystem that microbial communities having more cohesion (a metric that measures the complexity/interconnectedness of a given community) are more likely to be affected by homogenizing selection, and less cohesion are more susceptible to dispersal [15]. The microbial body size was revealed to structure the microbial community assembly in soil [16]. These findings bolster the likelihood of successfully applying established ecological theories and methods to understand the ‘rules’ of community assembly. However, the ecological study of plant microbiomes remains scarce; less attention has been given to elucidating how distinct ecological processes are operated and what the underlying ecological mechanisms are in structuring plant microbiomes. Additionally, most studies focused on the structure of the root-associated microbiota or bacteria; few have investigated bacterial and fungal communities simultaneously [17-19].

Woody plants provide multiple ecosystem services, e.g., climate regulation, nutrient cycling, carbon sink, production, and a reservoir of microbial diversity [20, 21]. Mulberry (*Morus* spp.) is a deciduous woody plant widely cultivated throughout subtropical and temperate regions [22], with broad applications in microbial biocontrol agents, medical engineering, and bioremediation [23-26]. In this work, we employed domesticated *Morus alba* L., as a model system, and recovered the bacterial (bacteriome) and fungal (mycobiome) communities from soil and plant niches (the episphere and endosphere of root, stem, and leaf, Fig. 1A), through amplicon sequencing of conserved phylogenetic markers. In view of the foregoing, we focused on 1) comprehensively assessing the biogeographic patterns of microbial communities (diversity, structure, co-occurrence network, species turnover, and assembly process) along the SPC and from episphere to endosphere (hereafter Epi–Endo); 2) exploring the potential ecological processes between the bacteriome and mycobiome across plant niches; and 3) underlying mechanisms that drive the patterning of plant microbiomes. We found that the bacteriome and mycobiome were strongly shaped by host selection, and more importantly, they had distinct biogeographic patterns within the soil-plant system, probably due to their different microbial phenotype (microbial size-dependent effect), as evidenced by ecological analysis and datasets from other plants. We thus explored this in more detail by proposing a “host selection-microbe size balance” hypothesis that smaller-size microbes were more likely to escape/disperse from inside niches. This hypothesis was confirmed by the stochastic dispersal experiments in which microbiotas stochastically dispersing out of the endosphere had significantly smaller size than endophytes.

**Fig. 1.**
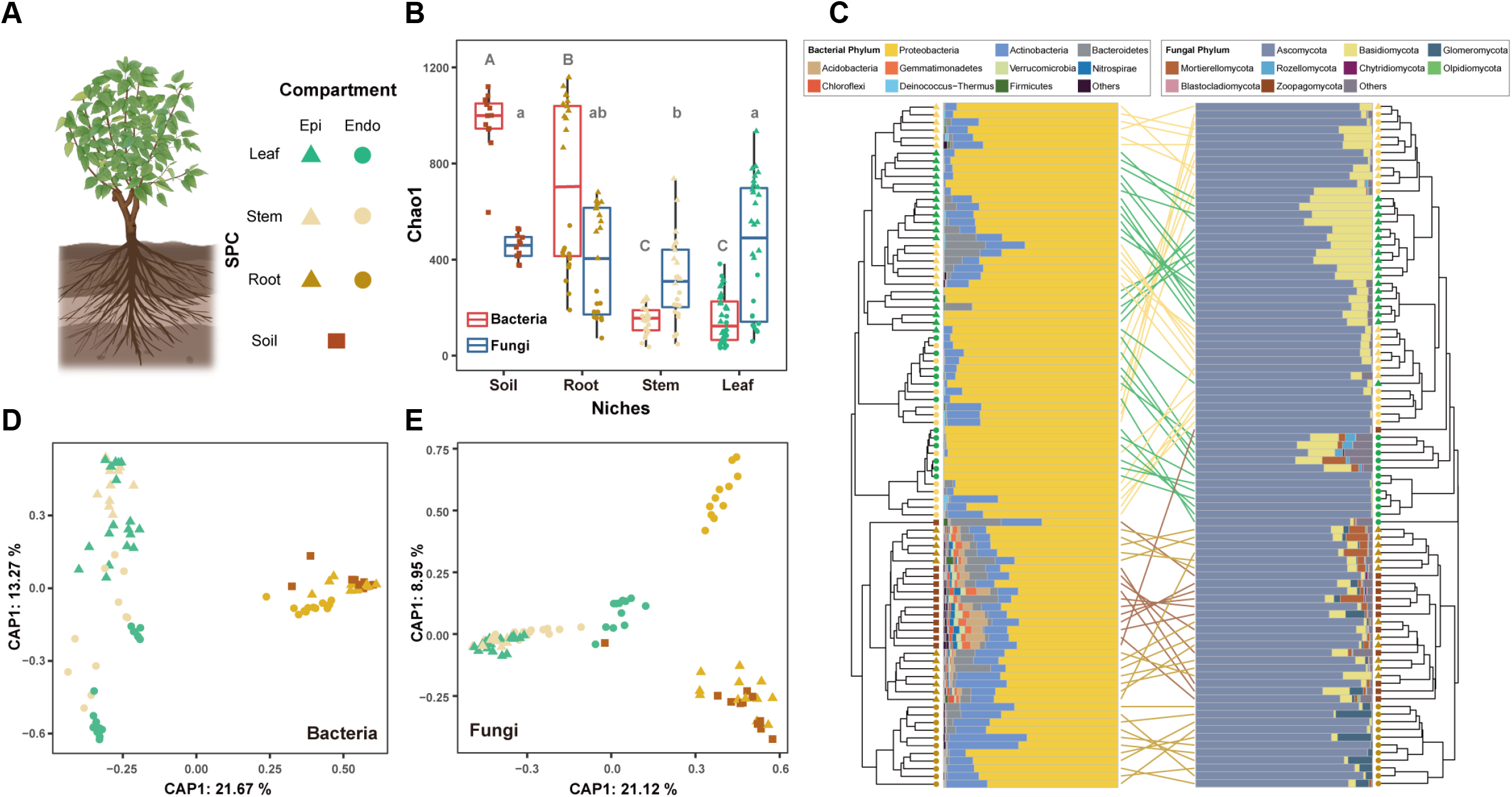
Host selection influences the diversity and taxonomic composition of bacteriome and mycobiome. **(A)** Schematic representation of seven niches amongst the plant-soil system in this study: soil, and the episphere and endosphere of root, stem, and leaf. Different colours differentiate between the fractions, soil (dark brown), root (brown), stem (light yellow), and leaf (green). Different shapes differentiate between epiphytic (triangle), endophytic (circle), and soil (square) communities. And the same shape and colour codes were used in the following figures. (**B**) The Tukey boxplot shows the diversity of bacterial and fungal communities in the soil, root, stem, and leaf. Each point represents one sample. Groups that do not share a letter are significantly different (*p* < 0.05); Boxplots show the median (line), 25^th^ and 75^th^ percentiles (box), and 1.5 × the interquartile range (IQR, whiskers). (**C**) Tanglegram visualization based on Bray-Curtis dissimilarities at the OTU level. Each node represents one sample. Bar plots show the taxonomic composition of bacteriome and mycobiome in each sample at the phylum level (bacteria, left panel; fungi, right panel). (**D, E**) Canonical analysis of Principal Coordinates (CAP) of bacterial and fungal communities across the plant niches. CAP ordinations using Bray–Curtis dissimilarity were constrained for the sample types. Axes report the proportions of total variation explained by the constrained axes.

## 2 Materials and methods

### 2.1 Sampling and data collection

*M. alba* L. plants were harvested in Hangzhou, Zhejiang province, China (30°18’6.31” N, 120°05’9.25” E). Samples in each niche (soil, the episphere and endosphere and root, stem, leaf) were collected from six individual plants in Dec. 2018 and Jun. 2019, placed on ice and shipped to the laboratory. The root samples were taken from the lateral roots of the plant after removing the loosely attached soil. Soils were sampled at 10 cm depth from the plant’s root and transferred to PowerBead Tubes (QIAGEN DNeasy PowerSoil Kit, Germany) as part of the soil niche. Epiphytic microbes were harvested from leaf, stem, and root samples, respectively, through extensive shaking in TE buffer supplemented with 0.1% Triton X-100 [27]. These washes were centrifuged at 8000 rpm for 10 min, and the pellet was dissolved in PowerBead Tubes as episphere fractions. Plant samples were then immersed in 0.25% sodium hypochlorite and 80% ethanol (3 min each) to remove the remaining microorganisms from plant surfaces and rinsed three times (1 min each) with sterile water. These surface-sterilized plant samples were used as endosphere fractions [28]. Overall, 96 samples: 12 soils (bulk soil), 24 root (rhizosphere and root endosphere), 24 stem (stem episphere and stem endosphere), and 36 leaf (phylloplane and leaf endosphere) samples, were obtained and transferred for DNA extraction and high-throughput sequencing of bacterial and fungal communities (96 × 2).

### 2.2 High-throughput sequencing

Samples were homogenized in a Precellys-24 tissue homogenizer (Bertin Technologies, France) at 5500 rpm for 45 s. Total DNA was extracted using QIAGEN DNeasy PowerSoil Kit according to the manufacturer’s instructions. The final DNA concentration and purity was determined by NanoDrop 2000 UV-vis spectrophotometer (Thermo Scientific, USA), and DNA quality was checked by 1% agarose gel electrophoresis. To avoid the influence of chloroplast DNA, a two-step PCR amplification protocol was used for bacteria detection [29]: The V4-V8 hypervariable regions of the bacterial 16S rRNA gene were first amplified with the primer set 799F-1392R; then the above PCR product was used as a template, and further amplified with the primer set 799F-1193R, which was designed for the V4-V7 hypervariable regions of 16S rRNA gene. The PCR reactions were performed in triplicate in a 20 μL mixture containing 4 μL of 5 × FastPfu Buffer, 2 μL of 2.5 mM dNTPs, 0.8 μL of each primer (5 μM), 0.4 μL of FastPfu Polymerase and 10 ng of template DNA. PCR reactions were conducted using the following program: initial DNA denaturation at 95 °C for 3 min, 27 (1^st^ PCR) or 13 (2^nd^ PCR) cycles of amplification with denaturation at 95 °C for 30 s, annealing at 55 °C for 30 s and elongation at 72 °C for 45 s, and a final extension at 72 °C for 10 min. For fungal detection, the fungal internal transcribed spacer ITS1 region was amplified using ITS1F-ITS2 primers under similar PCR reaction conditions with the annealing at 53 °C and the increased cycles to 35 [30]. Template-free water blanks were processed with the same DNA extraction and PCR amplification kits as negative controls to avoid reagent and laboratory contamination [31]. The PCR products were extracted from a 2% agarose gel and purified using AxyPrep DNA Gel Extraction Kit (Axygen Biosciences, USA) and quantified by QuantiFluor™-ST (Promega, USA) according to the manufacturer’s protocol. Purified PCR products were pooled in equimolar and sequenced (PE300) on an Illumina MiSeq platform (Illumina, USA) following the standard protocols provided by Majorbio Bio-Pharm Technology Co. Ltd. (Shanghai, China).

### 2.3 Sequence data processing

The acquired sequences were filtered for quality control as previously described [32]. Briefly, the resulting sequences after demultiplexing were merged using FLASH (v1.2.11) and quality filtered using fastp (v0.19.6) with the following criteria: (i) reads were truncated at any site receiving an average quality score <20 over a 50 bp sliding window. (ii) primers were exactly matched allowing 2 nucleotide mismatching, and reads containing ambiguous bases were removed. (iii) sequences whose overlap longer than 410 bp were merged according to their overlap sequence. Samples that failed to produce enough reads were excluded from the analysis. Any chimeric sequences were removed by UCHIME, and operational taxonomic units (OTUs) were clustered using UPARSE (v7.1) at 97% similarity cutoff [33]. The taxonomy of each sequence was checked using the RDP Classifier algorithm against the Silva (v132) and Unite (v7.2) databases under the condition of a 70% confidence threshold. Each OTU classification was screened, and singletons, Chloroplasts-assigned, and Mitochondria-assigned OTUs were removed for further analyses. OTU tables resampled to a minimum number of sequences from each sample (5532 for bacteria and 30324 for fungi). The desired sequencing depth was determined with rarefaction curves.

### 2.4 Co-occurrence relationship analysis

Network analysis was performed to explore co-occurrence patterns of bacteriome and mycobiome in each niche. The taxa with relative abundances above 0.01% were selected and those with low frequency (occur less than 25% in communities) were not included in the analysis [34]. A Spearman’s correlation between two OTUs was considered statistically robust if the Spearman’s correlation coefficient (r) was >0.6 and the *p*-value was <0.01 [35]. The resulting adjacency matrices were converted to network objects using the R package igraph (v1.2.6) and were visualized using the interactive Gephi platform. Network summary statistics and node centrality measures were calculated to compare network topologies (Node: number of used OTUs; Average degree: the extent of nodes connecting each other; Clustering coefficient: the ratio between the number of existing connections between a node and its neighbours, and the total possible connections; Average path length: Average of the shortest paths of any given node pair; Network Density: the ratio of the number of edges and the number of possible edges; Modularity: clustering degree in a network). The checkerboard score (C-score) was calculated to evaluate the real distributions for non-randomness of OTUs by examining the deviation of each observed metric from the average of the null model [36]. The values obtained were standardized to allow comparisons amongst communities using the standardized effect size (SES). Cohesion was calculated within both bacterial and fungal communities in each niche [37]. A negative cohesion was included since it was significantly related to community turnover and complexity. All negative null model-corrected correlations were averaged for each sample to obtain a connectedness matrix with average negative correlations. Finally, negative cohesion was calculated based on the connectedness and abundance of each taxon for a given community.

### 2.5 Microbial community assembly process analyses

Sloan’s neutral model analysis was performed to evaluate the contribution of neutral/niche-based processes in community assembly [38]. The model was adapted from the neutral theory and adjusted to fit large microbial populations. *R* square value represents the overall fit to the neutral model. The body size of each taxon was identified based on propagule size as previously documented [16]. Dispersal limitation represents those non-selection processes (e.g., heterogenous selection) contributing to more-dissimilar structures amongst communities, which could result from differential dispersal capacity or historical contingency (prior effects). To evaluate the dispersal limitation, the deviation in taxonomic diversity was calculated based on the Bray–Curtis dissimilarity-based Raup–Crick Index (RCI) for each taxonomic group. A mean RCI that is significantly greater or less than zero across all pairwise comparisons indicates greater (limited dispersal) or less (homogenizing dispersal) taxonomic turnover than the null expectation. RCI was calculated for each taxon detected in plant microbiome. To remove the influence of rare taxa, taxa with low prevalence (<50%) or low abundance (<0.5%) were excluded from correlation analysis. Alphaproteobacteria, Acidobacteria, Deltaproteobacteria, Bacteroidetes, Gemmatimonadetes, Verrucomicrobia, Chloroflexi, Firmicutes, Actinobacteria, and Gammaproteobacteria were analyzed in bacterial communities. While Chytridiomycota, Mortierellomycota, Agaricomycetes, Tremellomycetes, Cystobasidiomycetes, Microbotryomycetes, Leotiomycetes, Sordariomycetes, and Dothideomycetes were analyzed in fungal communities. Robustness regression (a form of regression analysis that is designed to minimize the violations of assumptions by the data-generating process) was performed to calculate the correlations between RCI and microbe size, due to considerable biological differences between taxa. Validation datasets (bacteriome and mycobiome of maize, wheat, barley, and Arabidopsis) were downloaded from NCBI [12, 28, 39]. The deviation in phylogenetic diversity was characterized by the β-nearest taxon index (βNTI) with the null model expectation generated with 999 randomizations [40, 41]. Weighted/unweighted size indexes were defined as the average size of all taxa detected in a community based on their abundance/presence.

### 2.6 Evaluation of stochastic dispersal

We carried out a stochastic dispersal experiment to evaluate the role of microbial body size. Seven wild mulberries were collected since they carry natural complex endophytic communities, which could be used as the pool of endophytic communities. Stems were surface-sterilized first with sterile water (two times, each with ultrasound sonification for 30 s), then in 8% Sodium hypochlorite (3 min), then in 70% ethanol (1 min) followed by washing in sterile water five times. To test the sterilization efficiency, 100 μL of wash water at the final step of surface sterilization was plated onto the LB media. Lack of bacterial growth from the last wash indicated the efficiency of surface sterilization. These endosphere fractions were transferred to 50 mL tubes with PBS, and were rotated 100 rpm using HulaMixer (Thermo Scientific, USA) to simulate the accelerated stochastic dispersal. Microbes stochastically escaping/dispersing from the endosphere at two time points (12 and 72 hours) were collected with additional five washes using sterile PBS. These washes were prefiltered through 11-μM pore size membranes to remove plant tissue debris, and then communities were collected on 0.22-μM pore size membranes. These membranes and remaining endosphere tissues were used for DNA extraction and amplicon sequencing to assess microbiota’s body size distribution.

### 2.6 Statistical analysis

Normal distribution and homoscedasticity were assessed by Shapiro–Wilk and Levene’s test, respectively. The α-diversity richness (Chao1) and evenness (Pielou) of each sample were calculated. The β-diversity was estimated based on Bray–Curtis distance (taxonomic) and UniFrac distance (phylogenetic). Constrained analysis of principal coordinates (CAP) was performed to evaluate the relationship between niches and community composition using the vegan package. Permutational multivariate analysis of variance was performed to disentangle variation in microbiota compositions using ADONIS in vegan. Differential abundance analyses were performed by DESeq2 to identify species enriched or depleted in each plant niche compared with that in soil. The potential functional profiling of bacterial and fungal communities was predicted using Phylogenetic Investigation of Communities by Reconstruction of Unobserved States [42]. The linear discriminant analysis effect size was used to identify biomarkers in different niches. Sample networks were constructed using phyloseq package. Molecular microbial source tracking was performed to assess the potential microbial source of the bacteriome and mycobiome in each niche using fast expectation-maximization microbial source tracking (FEAST) [43], and the Source Model of Plant Microbiome (SMPM) was constructed according to the sample network analysis and a previous study [12]. Given the high variability of the internal transcribed spacer region in fungi, we reconstructed a phylogenetic tree using the ghost-tree for validating the phylogeny-based metrics for fungal amplicon sequencing data [44].

## 3 Results

### 3.1 Host selection shapes the distinct structure and function of bacteriome and mycobiome

Alpha diversity analysis demonstrated that the diversity of both bacteriome and mycobiome was decreased from Epi–Endo (*p* < 0.001, Welch’s t-test) in each plant tissue (root, stem, and leaf); whereas the diversity along the SPC showed different patterns: the diversity of bacteriome decreased from soil to leaf, but mycobiome increased in leaf (Fig. 1B, Fig. S1). Notably, within belowground niches (soil and root), bacteriome has higher diversity than mycobiome; whereas mycobiome has a higher diversity within aboveground niches (stem and leaf). Mantel test showed that niches/samples sharing more closely related bacteriome harboured more similar mycobiome (*p* < 0.001, *R* =0.636, Bray-Curtis dissimilarity). Nonetheless, the tree obtained from bacteriome is not totally coherent with that from mycobiome (Fig. 1C), suggesting differential assembly processes between bacteriome and mycobiome in the soil-plant system. In addition, this pattern was still statistically significant by using Jaccard dissimilarity (*p* < 0.001, *R* =0.576, Mantel test) which does not incorporate information on relative abundance.

To unveil the variability in community structure between groups, Constrained Analysis of Principal Coordinates (CAP) was performed on bacteriome and mycobiome, respectively (Fig. 1D, E). The results showed that the bacteriome derived from the aboveground (leaf and stem) were grouped separately from those derived from the belowground (root and soil) (the first axis), while the variation of bacteriome between the episphere and endosphere was higher in aboveground niches compared with belowground niches (the second axis) (Fig. 1D). A similar clustering pattern was observed in mycobiomes on the first axis; however, the variation of mycobiome between episphere and endosphere was higher in belowground niches (Fig. 1E), suggesting complex factors in shaping the plant microbiome, such as the intense selection pressure in endophytic niches that reported previously [45, 46]. Therefore, it is crucial to study the epiphytes and endophytes (Epi–Endo) separately due to the contrasting host selection pressure.

Based on Bray–Curtis (taxonomy) and weighted UniFrac (phylogeny) dissimilarity, we next quantified the contribution of host effects, SPC and Epi–Endo, to community structure variations based on ADONIS test (Table 1). The taxonomic variation in bacteriome and mycobiome was mainly explained by SPC (*R*^2^ =0.291 and *R*^2^ =0.300, respectively), Epi–Endo (*R*^2^ =0.116 and *R*^2^ =0.069, respectively) and their interaction effect (*R*^2^ =0.099 and *R*^2^ =0.092, respectively), with slight effects of sampling time. Similar results were also observed in weighted UniFrac dissimilarity-based analysis and analyses at the phylum level (Table 1, Fig. S2). Moreover, the factor Epi–Endo affected bacteriome with a higher significance using weighted UniFrac (*R*^2^ =0.170) compared with that by Bray–Curtis (*R*^2^ =0.116); by contrast, the SPC showed more pronounced effects on mycobiome in weighted UniFrac (*R*^2^ =0.373) than in Bray– Curtis (*R*^2^ =0.300). We further examined the alterations in microbiome functions in plant niches. Functional potentials in both bacteriome and mycobiome were significantly clustered (*p* < 0.001, ADONIS) along the SPC (Fig. S3) and Epi–Endo (Fig. S4). A total of 15 functional pathways changed significantly between microbiomes in the episphere and endosphere, such as carbohydrate metabolism, cell motility, and membrane transport (Fig. S5).

### 3.2 Host selection shapes the patterns of bacterial and fungal co-occurrence networks

Plant microbiome assembly was further examined by investigating microbial co-occurrence network along the SPC and Epi–Endo (Fig. 2A, B). Results implied that microbial communities in plant tissues formed more complex associations (determined by clustering coefficients and graphic density than networks in soil, Fig. 2C, D) in responding to the host selection, and that induced network alterations may occur in a niche-dependent manner. Moreover, from Epi–Endo in each plant tissue, the bacteriome and mycobiome networks were switched to highly modularised but less connected (determined by modularity, average path length, and average degree, Fig. 2E, F). In contrast, fungal co-occurrence networks showed more sharply tuned alteration. As with network analysis, the C-scores showed higher standardized effect sizes (SES) in endosphere, which was more pronounced in mycobiomes than bacteriomes (Fig. S6, Fig. S7), confirming the effect of host plant on both bacterial and fungal endophytes, whereas fungi were more influenced. Moreover, negative cohesion was first applied to quantify the connectivity of bacteriome and mycobiome in each niche [15, 37]. Concordantly, negative cohesion gradually became stronger from the soil to plant niches except for mycobiome in the leaf endosphere (Fig. 2G, H). The negative cohesion in bacteriome was even stronger than that in mycobiome (*p* < 0.001, Student’s t-test). Remarkably, a drastic disparity between bacteriome and mycobiome was found when comparing epiphytes and endophytes. Compared with bacterial epiphytes, bacterial endophytes showed comparable or even significantly stronger cohesion in each plant tissue (*p* < 0.001, paired t-test, Fig. 2G). By contrast, mycobiome exhibited opposite patterns (*p* < 0.001, paired t-test, Fig. 2H), suggesting more susceptibility to homogeneous selection and dispersal in endosphere, respectively. Consistently, we found trends in homogenous selection in both the bacteriome and mycobiome (βNTI < 0, *p* < 1×10^−5^, Welch’s t-test, Fig. S8), with a stronger homogenous selection in plant niches compared with that in soil (*p* < 1×10^−5^, Welch’s t-test). Additionally, strengthened phylogenetic turnovers of endophytes than epiphytes were observed in bacteriomes but not in mycobiomes.

**Fig. 2.**
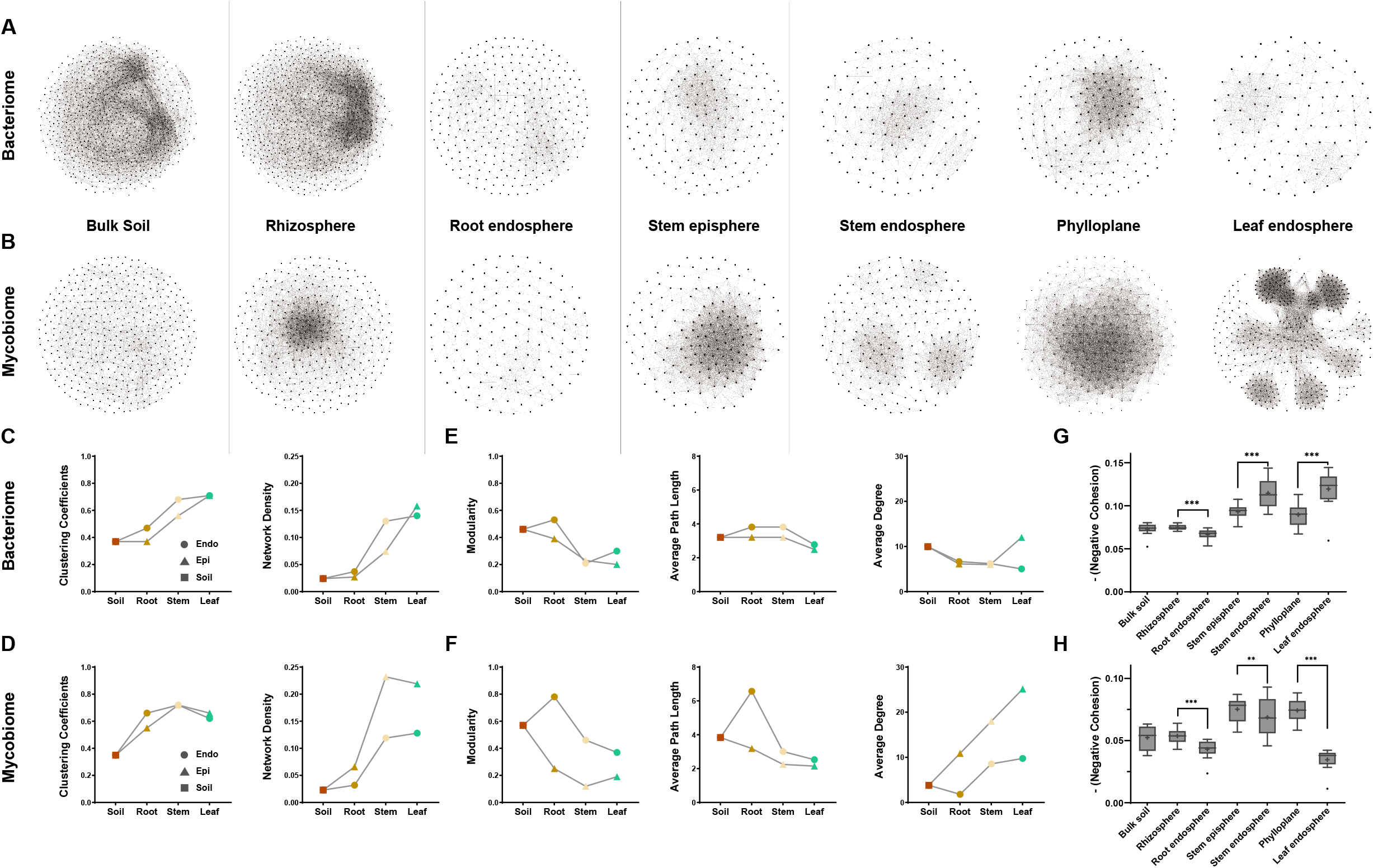
Host selection has remarkable effects on co-occurrence pattern across niches and cohesion strength may mediate the higher responses of mycobiome. Visualization of bacterial (**A**) and fungal (**B**) co-occurrence networks along the SPC and Epi-Endo. (**C**-**F**) The topological properties of bacterial and fungal co-occurrence networks, including clustering coefficients, network density, modularity, average path length, and average degree. Note that topological property of each co-occurrence network was significantly higher than that generated from their random networks with comparable parameters (*p* < 1×10^−5^, one sample z-test, 1000 Erdős–Rényi random networks), suggesting that networks are not random and that taxa involved in these potential associations are highly likely linked with each other. Changes in negative cohesion of bacterial (**G**) and fungal (**H**) community across niches. Stronger negative cohesion values were related to stronger complexity and alleviated community turnover.

### 3.3 Discrepancy in species turnovers between bacteria and fungi

Plant microbiome structure and co-occurrence network varied primarily across niches, indicating a general host selection. To further elucidate their assembly, we characterized the species turnovers by identifying the taxonomic groups associated with each niche. Several genera were significantly enriched in plant niches, such as *Pseudomonas* in the leaf, *Sphingomonas* in the stem, and *Steroidobacter* in the root (Fig. S9). Then we identified species clusters specifically enriched or depleted in plant niches compared with soil niches at the OTU level (Table S1). Totally, 9.3% of the OTUs (197 out of 2126 OTUs), mainly from the families Beijerinckiaceae (17 OTUs), Burkholderiaceae (16 OTUs), and Sphingomonadaceae (15 OTUs), were significantly enriched, while up to 25.8% of the OTUs (mainly from Nitrosomonadaceae) were depleted (Table S1). Similar patterns were also observed in mycobiomes. More importantly, our results revealed a right-skewed pattern of enrichment but a left-skewed pattern of depletion (Fig. 3). In other words, most enriched bacterial species were unique to each plant niches (albeit with some outliers in phylloplane and stem, Fig. 3A), whereas the depleted bacterial species were mainly identical amongst niches (more than half of the depleted OTUs, Fig. 3B). However, mycobiomes showed opposite patterns with left-skewed of enrichment and right-skewed of depletion, respectively (Fig. 3C, D).

**Fig. 3.**
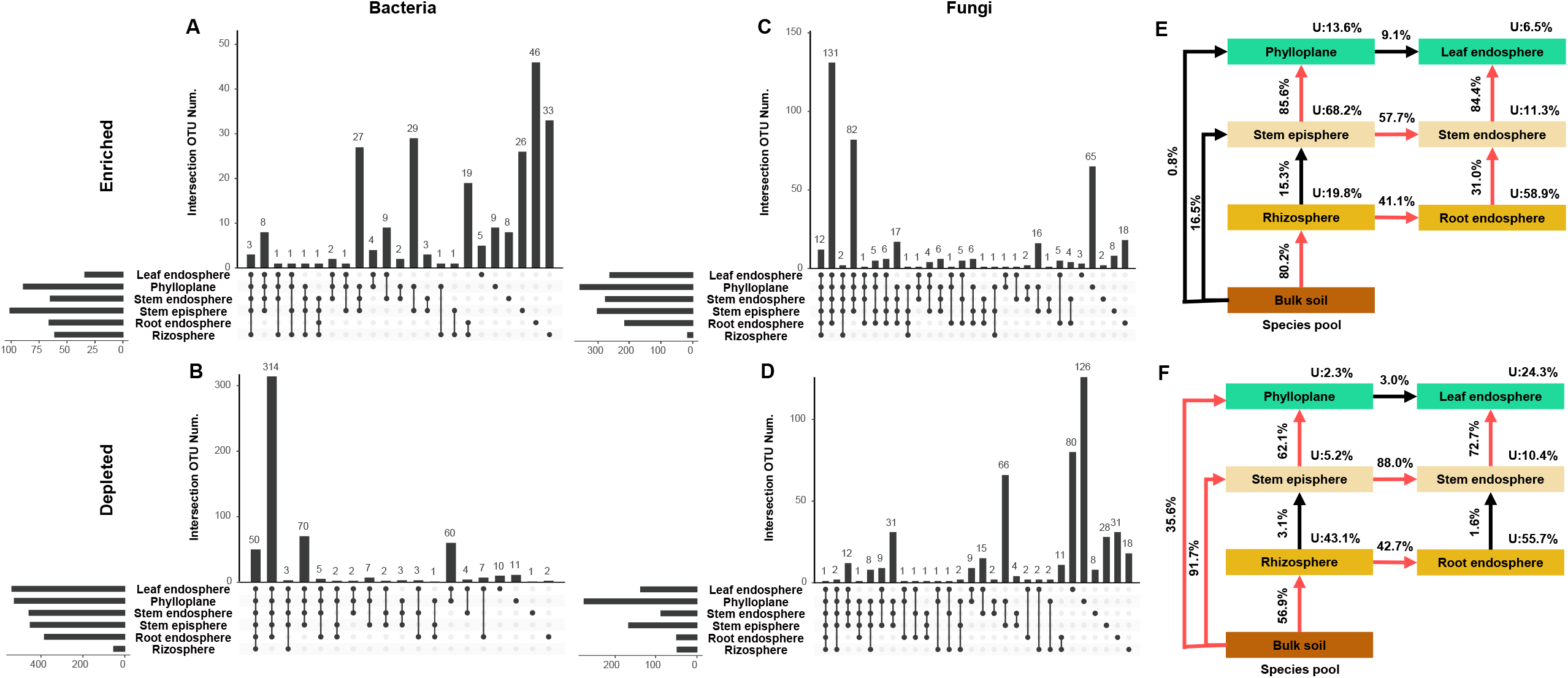
Enrichment and depletion in different plant niches shape distinct microbial species composition and show the discrepancy between bacterial and fungal species. Upset plots show the significantly enriched (**A, C**) and depleted (**B, D**) OTUs in different niches. DESeq2 was used to identify the differentially abundant taxa in each plant niche (adjusted *p* < 0.05, BH correction). Note that the distributions of enriched and depleted bacterial species are different from that of fungal species. (**E, F**) The microbial source model of plant microbiome (SMPM) of bacteriome and mycobiome. Colours differentiate four fractions, soil (dark brown), root (brown), stem (light yellow), and leaf (green). Lines with contributions larger than 30% were marked with red. U, unknown source.

Given the association between microbial communities amongst niches, we constructed sample networks of bacteriome and mycobiome, respectively (Fig. S10). Concordant with the CAP results, the soil microbiomes connected plant-associated microbiomes through root episphere following with root endosphere (Fig. S10), implying the potential routes of transmission and supporting the hypothesis that the microbiomes across plant niches come from the nearby species pool (niche). To further quantify the routes of transmission, the source model of plant microbiome (SMPM) was constructed and demonstrated that both the bacterial and fungal communities were mainly derived from bulk soils and filtered in plant niches. Specifically, for both bacteriome and mycobiome, the majority of species in leaf endosphere, phylloplane, stem endosphere, and rhizosphere, were potentially acquired from a nearby species pool (Fig. 3). The neighbour niches accounted for a smaller proportion of bacteriomes in stem episphere than that in phylloplane, indicating a potential contribution to stem episphere by other environmental sources (Fig. 3E). By contrast, most mycobiome species in the stem episphere were potentially selected from a nearby species pool with a higher known source value (94.8%) than that in the bacteriome (31.8%), and bulk soils were the main potential sources of stem episphere (Fig. 3F). The high contribution from bulk soil was observed in phylloplane mycobiome as well.

### 3.4 Size-dependent dispersal limitation in plant microbiota assembly

To further characterize assembly processes in shaping bacteriome and mycobiome, we applied several ecological approaches. Sloan’s neutral model was used to determine the relative importance of neutral processes in each niche. Consistent with cohesion and SMPM, fungal endophytes showed low values of fitness in plant tissues (0, 0.25, and 0.02 in root, stem, and leaf, respectively) (Fig. 4A, B), confirming that the dispersal of fungi could be constrained in the endosphere (fungi may hardly transmit in endosphere).

**Fig. 4.**
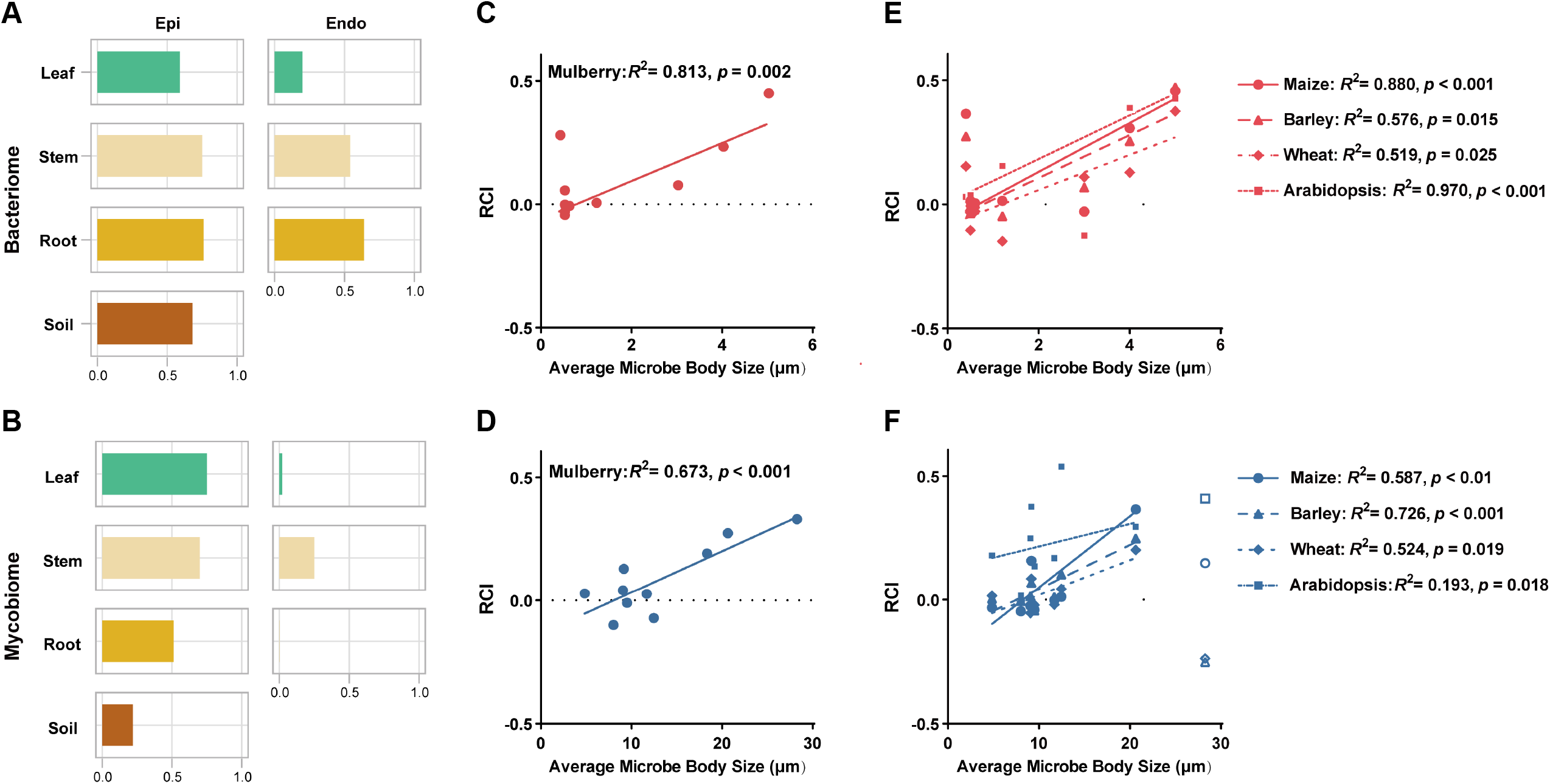
Size-dependent dispersal limitation in plant microbiome assembly. (**A, B**) The fitness of neutral model of the bacterial and fungal communities across niches. (**C, D**) The relationship between dispersal and organism body size of taxa within bacteriome and mycobiome. Each point represents a taxonomic group (with >50% prevalence or >0.5% relative abundance). A mean RCI (Bray–Curtis dissimilarity-based Raup–Crick Index, RCI) that is significantly greater or less than zero across all pairwise comparisons indicates greater (limited dispersal) or less (homogenizing dispersal) taxonomic turnover than the null expectation. (**E, F**) The relationship between dispersal and organism body size of taxa within bacteriome and mycobiome using four independent datasets of the microbiomes of four plants. Line types and point shapes represent taxa in different plants. Robust Linear Regression.

The results presented above showed that each community contained a majority of distinct organisms but also shared a proportion of similar organisms across niches, indicating the effects of host selection and dispersal on microbial community composition. This, together with contrasting patterns between bacteriome and mycobiome, prompted us to ask what factors drove the difference in assembly processes. One of the most significant differences in biology between bacteria and fungi is body size. Thus, we hypothesized that plant-associated microorganisms with larger sizes would be more restricted in their dispersal. We tested the size effects by calculating the correlations between body sizes and dispersal limitation of taxa within bacteriome and mycobiome, respectively (inferred by Raup–Crick Index, RCI). Most taxa showed limited dispersal (RCI > 0, Fig. 4C, D). Moreover, RCI was increased with larger body size in both bacterial and fungal taxa (*p* = 0.002, *p* < 0.001, in bacteriome and mycobiome, respectively, Robust Linear Regression), indicating that the limited dispersal is shaped in a size-dependent manner, albeit with small-size Alphaproteobacteria having a high RCI value (0.274). Additionally, concordant trends were observed between aboveground and belowground parts (Fig. S11).

To determine whether the *in-planta* size-dependent dispersal limitation acts as a general role in other plants, we analyzed multiple independent datasets in herbs, including maize, barley, wheat and *Arabidopsis thaliana*. The consistent trends were observed in bacteriome and mycobiome amongst these herb plants (Fig. 4E, F), albeit with the large-size Dothideomycetes having a low RCI value. As with our data, Alphaproteobacteria also had disproportional RCI values.

### 3.5 Balance between host selection and microbe size effect mediates plant microbiota assembly

Since statistical analyses denoted the significant underlying size-dependent effect in plant microbiome assembly, we further illustrated its relative importance and specific mechanisms. Previous studies have reported that the plant microbiome was determined by the orders of individual strain transferring, i.e., prior effects (prior microbes could inhibit through niche pre-emption or modification) [47]. Thus, we first speculated that the size effect should result in the statistically increasing size distribution from outside niches to inside niches as described in H_1_ (Fig. 5). Surprisingly, opposite patterns were observed in both bacteriome and mycobiome, validated by other plants (Fig. 6A, Fig. S12). Similar results were observed in plant niches along the SPC, particularly in bacteriomes (Fig. S13).

**Fig. 5.**
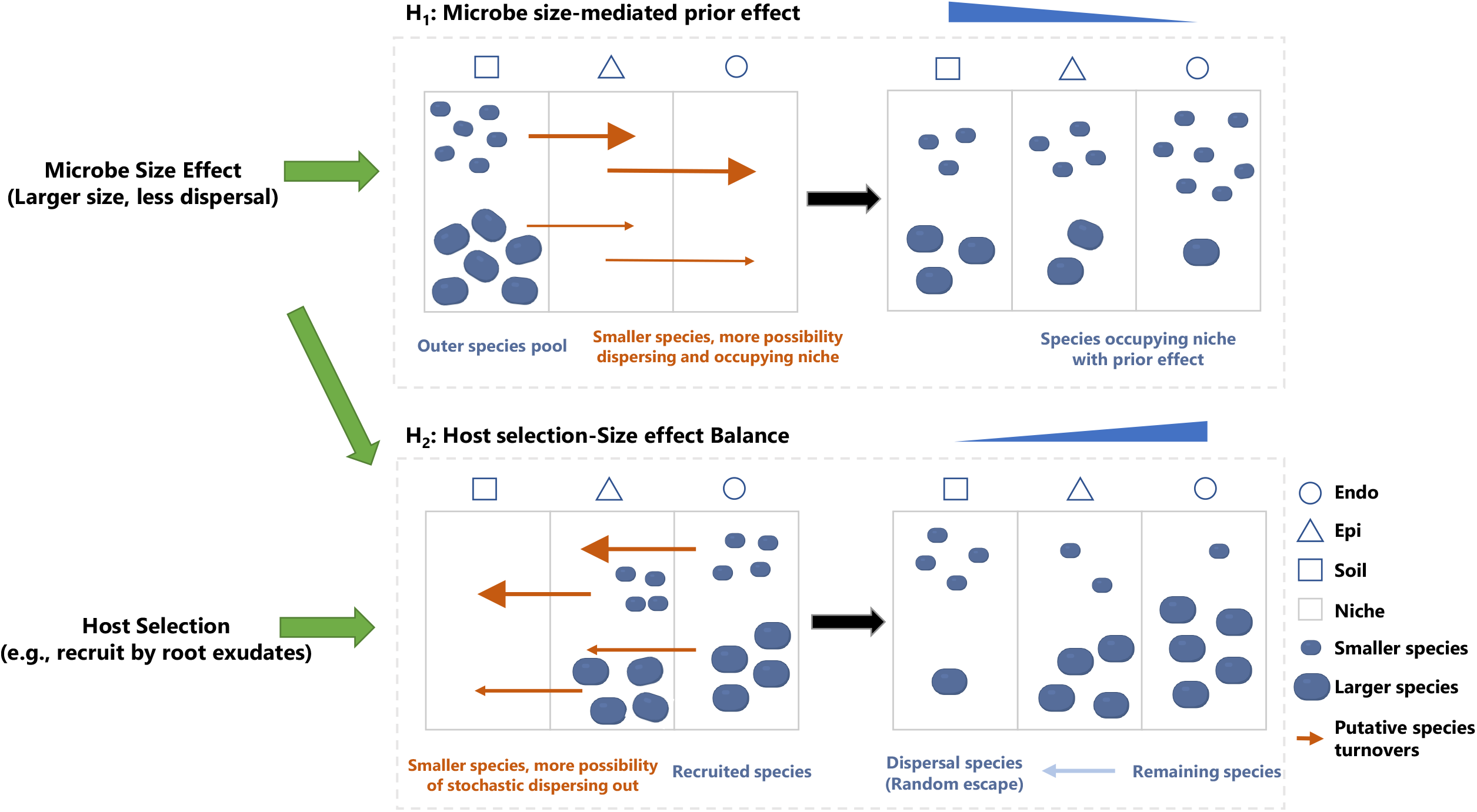
Schematic representation of the models. First model (**H**_**1**_) describes the microbial consortia assembled solely based on size effect through priority of smaller-size microbes. Limited dispersal of large-sized could result in less possibility of occupying inside niches (e.g., endosphere compared to episphere, and episphere compared to soil in belowground parts) during colonization due to prior effect (left panel in the first box). Thus, there would be decreasing size distribution from outside niches to inside niches (e.g., average microbial sizes decreasing from soil to episphere to endosphere) (right panel in the first box). Second model (**H**_**2**_) describes the microbial consortia assembled based on a balance between host selection and size effect. Host selection historically recruits and shapes the plant microbiome, and the microbe size affects the subsequent turnovers of the plant microbiome based on limited dispersal of larger-size microbe (left panel in the second box). In this case, there would be increasing size distribution from outside niches to inside niches (right panel in the second box.

**Fig. 6.**
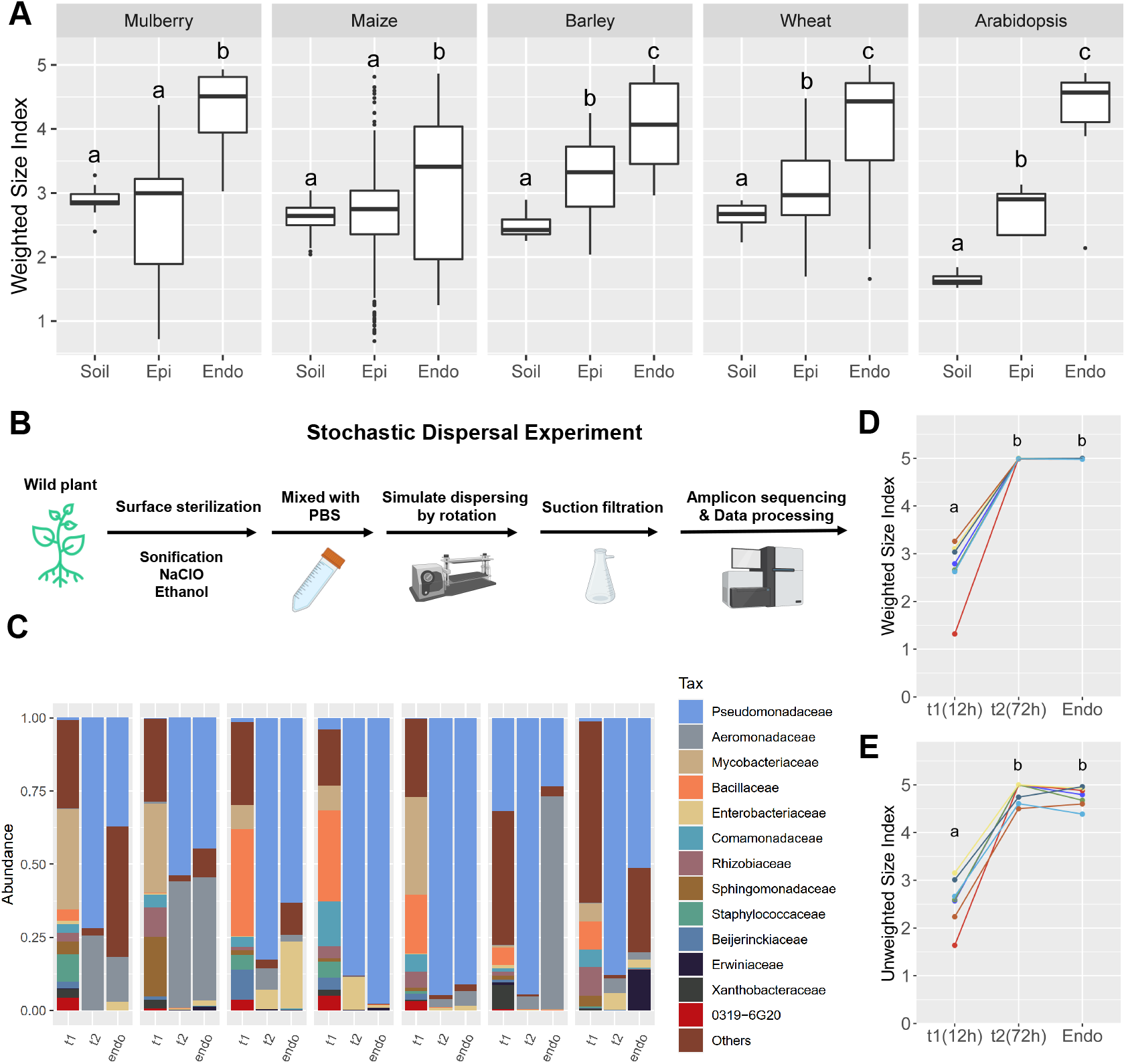
Balance between host selection and microbe size effect mediates plant microbiota assembly. (**A**) Boxplot of the weighted size index of bacterial communities across three niches (soil, episphere, and endosphere) in different plants including, mulberry, maize, barley, wheat, and Arabidopsis. Note that the distribution of microbe sizes across three niches disagrees with the model H_1_. (**B**) Stochastic dispersal experiment to determine the effects of stochastic dispersal/escaping by simulating the dynamic change of microbiota assembly (e.g., rainwater flowing over plant tissues). Briefly, wild mulberry plants were surface-sterilized; Endophytes were obtained through repeated shaking and rotating endosphere fraction (12 h and 72 h) in PBS. PBS solutions containing microbes (stochastically but extensively) dispersing out of endosphere and remaining endophytes were used for community characterization. Note that live microorganisms were detected from solutions based on an overnight culture on LB agar plates, suggesting that this developed procedure might provide a promising strategy to obtain endophytic community and to isolate endophytes. (**C**) Bar plots show the taxonomic composition of bacterial communities from seven mulberry plants used in stochastic dispersal experiment. The microbes dispersing during the first 12h (t1) and following 60 h (t2) and endophytic microbes are shown. (**D**) The weighted size index and (**E**) unweighted size index between t1, t2, and endophytic communities. Weighted/unweighted size index indicate the average size of all taxa detected in a community based on their abundance/presence. Groups that do not share a letter are significantly different (p < 0.05).

To reconcile these opposing results, we introduced the other factor, the host selection effect, into the model (H_2_, Fig. 5) since host selection could exert dominant effects in a short timeframe (historically affecting subsequent microbiome). For example, after inoculating cell mixture into the microbe-free plant, different compositions of endophytes could be observed at day0 [47], of which the timeframes are shorter than those of previously reported experiments at the week and month level [47-49]. We thus explored this in more detail by inferring that host selection might first recruit plant microbiomes, subsequently tuned by the size effect. In other words, smaller-size microbes were more likely to escape/disperse from inside niches, and thus larger-size microbes could occupy these fleeting niches, contributing to larger sizes in inside niches.

We designed a stochastic dispersal experiment using plants without epiphytes to determine the effects of stochastic dispersal/escaping by simulating the dynamic change of microbiota assembly (Fig. 6B, details see “Methods”). Because the size effect is independent of species identity and bacteriome showed more explicit patterns (possibly due to their higher inter-taxon size variation and simpler cell structure), we characterized the bacterial community to test the H_2_ (Fig. 6C). The microbes dispersing during the first 12h (t1) had significantly altered size distribution compared with those in the following 60h (t2) and endophytes (Fig. 6D, E), where large-size taxa were primarily detected, such as Pseudomonadaceae and Aeromonadaceae (Gammaproteobacteria). No significant difference was observed between t2 community and endophytes in microbe sizes (*p* = 0.350) and community composition (*p* = 0.373, ADONIS), possibly due to the low resolution of taxonomic sizes (documented at phylum and class level). These results confirmed that the smaller-size microbes were more likely to disperse out of the endosphere.

## 4 Discussion

Deciphering the principles governing the plant microbiome assembly can advance our mechanistic understanding of plant microbiome manipulation to promote plant performance [50, 51]. Assembly of plant microbiomes is mainly determined by niche-specific selection with marginal effects such as cultivar, age, and fertilization [52, 53]. Our data supported the dominant role of host effect (niche-specific selection, accounting for 50.6% and 46.1% in bacteriomes and mycobiomes, respectively), with a relatively minor role of the sampling time in regulating microbiomes in mulberry. Despite the distinct microbial communities across niches, they are not separate but closely linked, as evidenced by our results. Factors such as the provision of specialized nutrients amongst niches and immune systems [54, 55] could shape the distinct composition along the SPC (e.g., light, nutrients) and Epi-Endo (e.g., host immune system) [56, 57]. By rhizodeposition and root exudation, plants recruit and shape a distinct and diverse microbiome around the root [51, 58, 59]. Root epiphytes can become endophytic (e.g., through endodermis and pericycle, reaching the xylem vessels) and be transferred internally to the aboveground tissues, and endophytes can also arise from ingression into leaf endosphere following epiphytes colonization [60, 61]. Then, factors driven by the host generate a more distinct endophytic community dependent on the plant–microbe interactions [62]. For example, bacteria such as host-enriched Burkholderiaceae and Sphingomonadaceae have also been reported enriched in other woody plants [63, 64]. Therefore, not only the soil–plant (SPC) but also the episphere–endosphere (Epi–Endo) are really parts of the continuum for plant microbiomes [60, 65] (Fig. 1A). Besides, the assembly process may be different between bacteriome and mycobiome, as shown in the source tracking analysis. A large proportion of mycobiome in aboveground parts is primarily acquired from the soil, possibly resulting from their different spread strategies (such as spores spread in the air). Nevertheless, the contributions of other potential sources (e.g., air, microbe-carrying animals in the field, and vertical transmission) are worth considering. A recent field study found that environmental sources contributed to phylloplane microbiomes by comparing maize and fake plant [66]. Similarly, Beijerinckiaceae, enriched in mulberry, are reported as core members in aerosol microbiomes [67]. On the other hand, plant microbiomes are also balanced through microbe–microbe interactions [68, 69], as shown in co-occurrence networks amongst niches. Despite sequentially decreased microbial diversity, the associations were more complex from soil to leaf, in contrast to a previous report on herbs [12]. This might be explained by the woody plant that we studied different from herbs in multiple facets. Woody plants produce lignin, grow slowly, and have a strong stem, longevity, and late sexual maturity, which makes their physiology, development, and ecology different from annual plants [20, 70], resulting in, for example, a relatively higher diversity of leaf microbiome.

Despite the major roles of compartmentalization in plant microbiome composition and network, the ecological patterns remain largely underexplored in bacteriome and mycobiome. Recent research has underlined the role of eco-evolutionary processes in the assembly of plant microbiomes, such as dispersal (movement of microorganisms between different niches, or historical contingency), selection (biotic and abiotic effects causing fitness differences), ecological drift (stochasticity), and diversification [54, 71, 72]. Our study offers another perspective based on the application of ecological analysis on complex and diverse microbiota composition rather than the limited number of microbial strains. To the best of our knowledge, our study, for the first time, revealed the size-dependent effect in regulating plant microbiome assembly. Bacteria have typically small size and display unicellular growth, allowing them to colonize xylem vessels and access other plant parts in the endosphere much easier [58, 73]. By contrast, translocation processes (dispersal in plant body) in fungal species in the endosphere may be restricted to some extent due to their larger size, as reported in soil [74] and marine [75] ecosystems. The assembly processes could also be affected by plant–microbe interactions in a specific niche. For instance, common symbionts, Alphaproteobacteria and Dothideomycetes, exhibited disproportionally high/low RCI despite their small/large body sizes. This could be ascribed to their close association with plants in specific niches, such as *Rhizobium* and *Azorhizobium* in root [76, 77]. Collectively, our findings support a general relationship between body size and ecological processes in plant microbiomes. More importantly, we showed that the size effect does not play roles alone but through a balance with host selection, as described by the H_2_. This could also partially explain their differences in enrichment and depletion patterns due to tuned host selection-size effect balance since fungi have larger size, greater niche overlap, and different sensitivity to biotic and abiotic factors [54, 60, 78]. Moreover, the developed stochastic dispersal experiment enables us to collect endophytes (even those alive) that disperse out of the endosphere in a stochastic way. Nonetheless, other microbial features such as shape and mobile capability may also exert effects, as reported in environmental media [79, 80]. In addition, mechanisms, such as the size-plasticity hypothesis, might also play roles. Further analytical developments and experimental studies are required to provide additional supportive evidence—for instance—using the microbe with manipulated size through a metabolic sensor or reducing FtsZ (a cell division protein) levels without impacting growth rate [81, 82].

Breeding toward improved ecological plant–microbiome interactions requires improved knowledge of ecological processes/principles underlying their patterns [83, 84]. Moreover, the assembly processes of plant microbiomes are essential not only for plant performance but also for connecting soil, plant, animal, and human. To sum up, this study constitutes first detailed investigation of *in-planta* biogeographic patterns of plant-associated bacteriome and mycobiome in mulberry. Based on ecological analyses and stochastic dispersal experiment, we showed the size-dependent mechanisms ruling the population of plant microbiomes, and how it together with balanced host selection works in shaping the plant microbiome. These findings shed light on the complex host–microbe interactions within the tree plant and may provide implications for developing rational ecological-based designs of plant-beneficial synthetic microbial consortia.

## Conflicts of interest

We declare no conflict of interest.

## Acknowledgements

This work was supported by grants from the National Natural Science Foundation of China (Grant No. 32022081 and 31970483), Zhejiang Provincial Natural Science Foundation of China (LZ22C170001), China Agriculture Research System (Grant No. CARS-18-ZJ0302), key laboratory of silkworm and bee resource utilization and innovation of Zhejiang Province (2020E10025) and the Max Planck Society, Germany. The funders had no role in study design, data collection and interpretation, or writing the manuscript. Schematic representations were generated with the support of Biorender (©BioRender - biorender.com).

## Data availability

The sequence data were deposited into the NCBI Sequence Read Archive (SRA) database BioProject (Accession Number: PRJNA549049). Scripts used in this study are available at the GitHub repository (https://github.com/kingtom2016/mulb_niche).

## Figure captions

**Table 1 The effect of plant niches on the taxonomic and phylogenetic structure of bacteriome and mycobiome.**

## Notes

### Competing Interest Statement

The authors have declared no competing interest.

